# Fundamental limits on fluorescence lifetime estimation using optical gating

**DOI:** 10.64898/2026.11.25.769309

**Authors:** Pingchuan Ma, Linghao Hu, Adam J. Bowman

## Abstract

Fluorescence lifetime imaging (FLIM) is a promising approach for quantitative measurement in biology. Traditional lifetime measurements have limited photon throughput and slow acquisition speed due to photon counting electronics and point-scanning detectors. Wide-field imaging with optical gating can overcome these limitations. We analyze lifetime estimation precision for both single- and bi-exponential decays with optical gating waveforms. We focus on electro-optic (EO-FLIM) gating approaches that can be implemented all-optically with no photon loss and compare them to standard binning methods. Fisher information (FI) is used to calculate the Cramér-Rao lower bound (CRLB) for decay parameter estimation for each gating strategy, which is then verified with maximum likelihood estimation on Monte Carlo simulated data. We present fundamental limits for both single and bi-exponential estimation with resonant, step, and square gating waveforms. For single exponential decays, a single step or resonant gate approaches the FI limits from time correlated single photon counting (TC-SPC) within a factor of 1.2-1.6. We map the parameter space for laser excitation waveform, gate waveform, and gate number. For bi-exponential estimation, at least 3 parallel step or resonant gates are required to estimate the full parameters. With prior knowledge of one lifetime, two optimally-chosen gates can approach the TC-SPC limit within 1.6 (step gate) or 2.9 (resonant gate). Estimation precision converges within 5-10 gates across conditions with one known lifetime component and within 10-15 gates with no prior information. Asymptotic performance with increasing gate number achieves between 3.1 (step) and 3.7 (resonant) the TC-SPC limit with one known lifetime and between 3.5 (step) and 8.8 (resonant) with no prior information. These results inform the design of sparse gating strategies that maximize throughput and per-photon information for lifetime estimation. Optical gating enables future experiments for high-throughput sensing and multiplexing of fluorescent biosensors without relying on time-resolved detector electronics.

## INTRODUCTION

Fluorescence lifetime microscopy measures the time between the excitation of a fluorophore and the emission of a fluorescence photon, which occurs within a few nanoseconds for fluorescent proteins and most chemical dyes used in biology (1). The lifetime reflects the intrinsic state of the fluorophore and can be used for sensing the local environment through processes that modulate non-radiative decay paths. Lifetime offers quantitative advantages compared to measuring changes in fluorescence intensity. It is independent of excitation intensity, probe expression levels, and photobleaching, allowing cellular signals to be tracked across both transient and chronic timescales as well as across model systems and animals. These advantages are being realized in applications ranging from the study of fast and slow neuronal signals (2–4) to imaging of metabolites (5, 6), intracellular signals (7–9), and physical environment (10–12). However despite its broad range of applications, lifetime imaging has been limited to low throughput acquisition that is often orders of magnitude slower than standard wide-field and intensity based approaches or, alternatively, to specialized image sensors that have high levels of measurement noise.

Traditional FLIM approaches such as time-correlated single photon counting (TC-SPC) sample photons using large number of picosecond time bins to construct fluorescence lifetime decay histograms (13) (Fig. 1a, left). While highly accurate, TC-SPC is limited in throughput and acquisition speed, generally requiring seconds or even minutes to acquire an image depending on field of view. This limitation is due to point scanning and the limited photon acquisition rates enabled by photon counting electronics. Emerging FLIM methods combine wide-field imaging with various temporal gating strategies to achieve fast fluorescence lifetime imaging. For example, single photon avalanche diode (SPAD) array cameras achieve scanless fluorescence lifetime imaging by employing electronic gating of photon detections within each camera pixel but discard photons arriving outside the gate window (14–16). Modulated cameras can also implement sinusoidal gates within each camera pixel (17, 18), but commercial implementations exhibit limited frame rate and approximately 30 times higher read noise than scientific CMOS cameras(19). We have recently developed electro-optic fluorescence lifetime imaging microscopy (EO-FLIM) to achieve low-noise and high photon efficiency fluorescence lifetime measurements in wide-field imaging (2, 20–22). EO-FLIM uses electro-optical crystals as polarization gates to switch fluorescence photons into gated (G) and ungated (U) paths after a polarizing beam-splitter, both of which are either captured directly by low noise scientific cameras (Fig. 1a, right) or passed into additional crystals for series gating (Fig. 1c). Shot noise in each final detected intensity channel limits estimation precision. Because low-noise and high quantum efficiency detectors such as back-thinned scientific cameras can be used, this approach has many metrological advantages. It has demonstrated kilohertz imaging of neuronal voltage in vivo with subthreshold resolution, rejection of noise and motion artifacts, and picosecond level pixel noise across FLIM images (2, 22) as well as applications in single-molecule, super-resolution, and light sheet microscopy(21, 23, 24).

**Figure 1:**
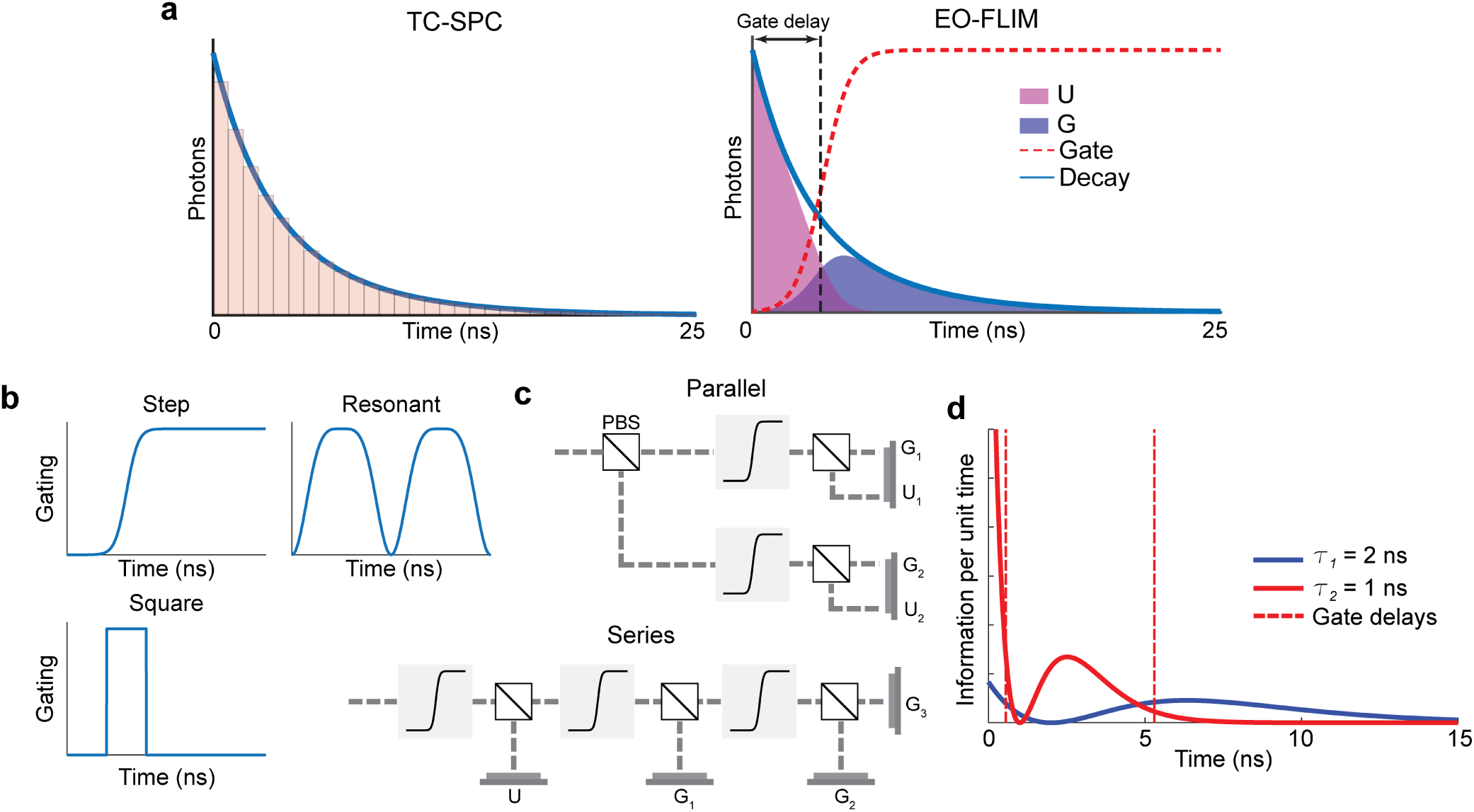
Fluorescence lifetime gating strategies. **a)** Conventional time-correlated single photon counting (TC-SPC) relies on many narrow time bins to build a histogram describing photon arrival times. EO-FLIM uses a pair of lossless complementary gates which may have a gradual rising edge imposed by drive electronics for an electro-optic crystal. **b)** Several gate functions (step, resonant, or square shapes) can be realized for EO-FLIM. **c)** Each EO-FLIM gate here is implemented by an electro-optic polarization modulator (shaded box) followed by a polarizing beam splitter (PBS). EO-FLIM gates may be combined in parallel or in series to realize multiple gates on an array of intensity detectors. **d)** The temporal distribution of information density (see section S1.4 in the Supporting Material) is plotted (information per unit time) for two lifetime components in a double exponential fluorescence lifetime decay (*τ*_1_ = 2 ns, *τ*_2_ = 1 ns). Dashed line shows the two optimal step function gate delays to estimate *τ*_1_ with known *τ*_2_ (obtained from Fig. 3)

Applications of FLIM rely on reading out spatial or temporal changes in the nanosecond lifetime, which may generally follow a single or multi-exponential process. Single exponential readout can use a mean or intensity-weighted average lifetime to report a continuous underlying variable. In other cases, it is valuable to have the capability to measure and unmix two or more exponential decay components. For example, resolving two decay components can derive substrate bound and unbound fractions of a FRET biosensor reporting kinase activity (7, 25), measure two autofluorescence decay components for optical metabolic imaging (26, 27), or unmix overlapping probes within a single spectral channel (28–31).

In TC-SPC, lifetime components are generally estimated by fitting finely-sampled decay histograms to single or multiple exponential equations (8, 13). Bin widths are usually in the 10s of picoseconds and hundreds or thousands of time bins are acquired. The lifetime estimation limits of TC-SPC have been evaluated by both analytical (32–36) and numerical (28, 36–40) methods. However, alternative gate strategies have only been analyzed in limited cases such as traditional frequency domain FLIM (41) and related in-camera modulated pixels (17, 42) or imaging with gated optical intensifiers using sequential or lossy gating protocols (43–46). Previous analyses have been limited to the contexts of lossy gates (e.g. gated intensifiers) or in-pixel gating where electronic parameters and pixel noise often dominate, and they have not evaluated gradual gating waveforms and lossless gating strategies such as those possible with EO-FLIM. Further complicating the problem is the large parameter space for experimentally relevant EO-FLIM setups: **1)** EO-FLIM is applicable to different excitation laser waveforms such as pulsed, sinusoidal, and square excitation which involve large potential cost and excitation power trade-offs. **2)** Multiple types of gate functions can be experimentally realized with electro-optic approaches including step, resonant, and square gating waveforms (Fig. 1b). **3)** The delay between laser excitation and gate application can be arbitrarily tuned (Fig. 1a). **4)** Various numbers of parallel or series gates, each with arbitrarily tuned delays, can be applied (Fig. 1c). This flexibility allows EO-FLIM to be applied in many experimental contexts and with diverse fluorescent probes, but it also requires systematic evaluation and experimental design to achieve optimal lifetime estimation performance.

In the following work, we analyze step and resonant gates in parallel configurations where photons are evenly divided among a number of independent gated measurements. These gates may be acquired truly in parallel by optical splitting with beam-splitters (Fig. 1c, top) or by dividing photons among camera exposures (i.e. a separate exposure for each gate delay). These are mathematically equivalent. Optical gating using series EO-FLIM units (modulator and polarizing beam-splitter) is also possible (Fig. 1c, bottom) and can be used to create square gates where we expect the best estimation performance i.e. at each time a photon can only be routed onto one intensity detector. In practice, the series optical arrangement can realize square gates from step voltage waveforms which may have practical advantages for experimental implementation. The resonant gating waveform we analyze takes the form 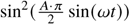, which corresponds to the transmission of a birefringent phase modulator driven with a sinusoidal voltage of amplitude *A* (units of *V_π_*, the half-wave voltage) between crossed polarizers (22).

Fisher information theory (32, 36, 47, 48) provides a framework to systematically evaluate the achievable per-photon information in fluorescence lifetime measurements. The temporal gates in EO-FLIM must optimally interact with the decay over time to enable efficient information extraction for lifetime estimation (Fig. 1d). Inspired by this, here we develop a computational framework based on Fisher information analysis to evaluate lifetime estimation in single and bi-exponential decays across different optical gating schemes and varying modes of laser illumination. Fluorescence lifetime estimation precision *σ_τ_* is generally given by 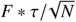 where for an ideal shot-noise-limited measurement with photon counting *F* = 1. The F number provides a quantitative method to compare gating approaches and FLIM modalities. We calculate the Cramér–Rao lower bounds (CRLBs) and the corresponding F numbers to evaluate the best achievable estimation performance for different gating schemes. We also use Monte Carlo simulation and maximum likelihood estimation to independently verify these results. For each gating scenario, we provide comparisons to ideal single-photon counting using 256 time bins distributed across a 25 ns (40 MHz) laser period as the gold standard method. Our gating scenarios reflect shot-noise limited measurements that may be achieved using low noise scientific camera sensors. We also explore the effect of gate number and show that in many cases fundamental information bounds can be approached using less than 10 gates in simple experimental designs. This computational framework enables optimal design of gating strategies to achieve the best per-photon lifetime estimation in future optically gated FLIM experiments.

## METHODS

This section describes Fisher information analysis of Cramér–Rao lower bounds (CRLBs) for decay parameter estimation as well as cross-verification with maximum likelihood estimation (MLE) using simulated measurements. The Fisher information analysis was adapted from previous studies on TC-SPC (32) to the case of EO-FLIM with distributed optical gating waveforms and complementary lossless gates as described below and in the Supporting Material.

### 0.1 Exponential Fluorescence Lifetime Decay Models

For an instantaneous excitation pulse at *t* = 0, a bi-exponential fluorescence lifetime decay is described by:

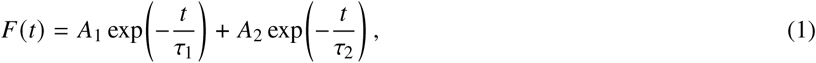

where *F*(*t*) is the fluorescence intensity and *A_i_* is the initial intensity of the component with lifetime *τ_i_* immediately after excitation. FLIM systems use excitation with period *T*. In this study we analyzed cases with 40 MHz laser repetition rate and 25 ns period. In the steady state, the fluorescence observed within one period (0 ≤ *t* < *T*) also contains the tails of all preceding pulses:

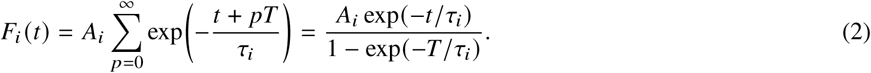

Integrating over one period gives the total photons as 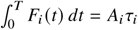. Normalizing *F_i_*(*t*) then gives the photon probability density for this component:

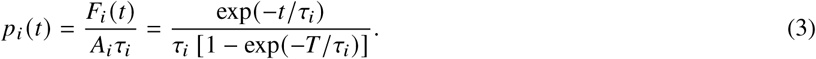

We numerically implemented the photon probability density with K time bins within excitation period T with bin width Δ*t* = *T*/*K* (*K* = 100000; T = 25 ns). The probability that a photon from component *i* falls into the *k*^th^ bin is

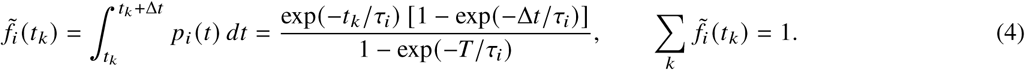

The expected photon count at time *t_k_* for a bi-exponential decay is then

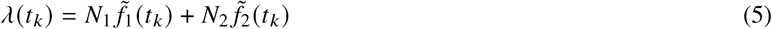

where *N_i_* ∝ *A_i_τ_i_* is the expected total number of detected photons from lifetime component *i*.

In EO-FLIM, each parallel gate will generate one set of complementary gated (*G*) and ungated (*U*) measurements that sum to the total intensity going into a given beam splitter (Fig. 1a, right). In the case of series gates there are not complementary pairs, but only a single U channel depending on the configuration (Fig. 1c, bottom). The total number of paired parallel measurements will be twice the number (*M*) of gates deployed. To generalize the *G* and *U* measurements, we define each expected value in gate *m* (*m* = 1, 2, …, *M*) as:

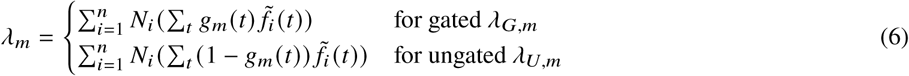

Here, *g_m_*(*t*) represents each of the gating functions applied (i.e. transmission from 0 to 1 — see Section S1.2 in the Supporting Material for detailed definitions).

### 0.2 Fisher Information Matrix

The Fisher information matrix is constructed from the derivatives of the measurements with respect to each of the decay parameters ***θ*** = [*τ*_1_, *N*_1_, *τ*_2_, *N*_2_] for a bi-exponential decay. Previous studies have established the matrix elements for TC-SPC measurements (32). We modified this matrix to reflect measurements generated by temporal gates in EO-FLIM. The derivatives of the gated measurements *λ_m_* with respect to the photon numbers *N_i_* of each lifetime component are defined as *J_i,m_*:

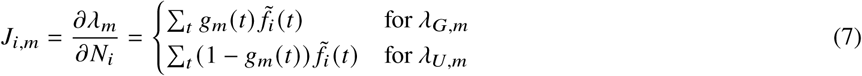

The derivatives of the measurements with respect to the lifetimes *τ_i_* are defined as:

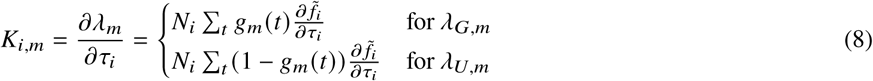

where detailed calculation for 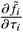 is described in Section S1.1 in the Supporting Material. The Fisher information matrix *I* for a bi-exponential decay parameter vector ***θ*** = [*τ*_1_, *N*_1_, *τ*_2_, *N*_2_] is constructed as:

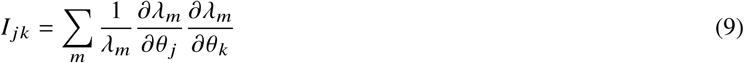

where *I* is a 4×4 matrix and the elements are calculated from paired *J* and *K* terms (see Section S1.3 in the Supporting Material).

### 0.3 Cramér-Rao Lower Bound

The Cramér-Rao lower bound (CRLB) for each parameter is obtained from the diagonal elements of the inverse Fisher information matrix:

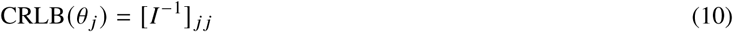

The CRLBs establish the theoretical lower bounds on the variance of parameter *θ_j_* estimation using any unbiased estimator. For parameters that are known and not estimated from the data, the corresponding rows and columns are removed from *I* before inversion. The CRLB is then directly related to the F number in FLIM estimation as the photon normalized relative precision for each parameter:

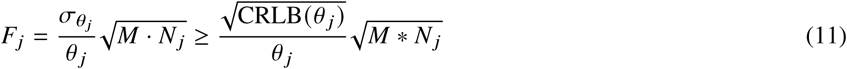

Where M accounts for equal photon division among parallel gates (step and resonant). For series square gating M = 1 here.

### 0.4 Maximum likelihood estimation (MLE) from EO-FLIM Measurements

EO-FLIM measurements consist of gated (*G_m_*) and ungated (*U_m_*) photon counts for each gate function (*m* = 1, …, *M*). Each of the measurements are modeled as independent Poisson random variables with means *λ_G,m_*(***θ***) and *λ_U,m_*(***θ***). The likelihood of observing the full set of measurements is therefore the product of the individual Poisson probability distributions for each intensity measurement:

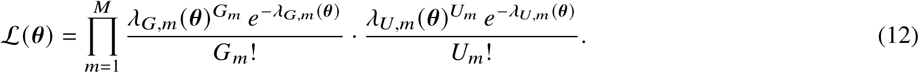

Taking the logarithm and dropping the terms ln *G_m_*! and ln *U_m_*!, which do not depend on ***θ***, gives the log-likelihood:

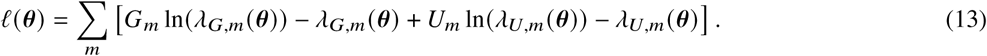

The parameters ***θ*** (*τ*_1_, *τ*_2_, *N*_1_, *N*_2_) are estimated by minimizing the ℓ(***θ***). To check the CRLBs derived from the Fisher information, we generated *G_m_* and *U_m_* in repeated Monte Carlo simulation trials and estimated the decay parameters from each trial by MLE. The F number was then calculated from the standard deviation of the estimates across trials and compared with the value predicted from the Fisher information (see Section S3 in the Supporting Material).

## RESULTS

### 0.5 Single exponential decays

We first performed a comprehensive analysis for gating-based estimation of single exponential lifetimes, exploring three gate functions (step, resonant, and square), three laser illumination waveforms (pulsed, sinusoidal, and square), and varying the numbers of parallel or series gates applied to the fluorescence photons.

A single gate with two complementary measurements (G and U) is sufficient to estimate single exponential lifetime (Fig. 2) with F = 1.24 (step) or 1.48 (resonant), which matches previous analysis (47). We calculated CRLBs to derive the F numbers of lifetime estimation using a single gate function across the parameter space (Fig. 2a-d and Fig. S1). The F numbers using the optimal gates across different lifetimes are higher than TC-SPC for step and resonant gates (Fig. 2c) by factors of 1.2 and 1.6 respectively. Note that an equivalent measurement precision to TC-SPC would require *F*^2^ more photons, but camera-based detection provides several orders of magnitude in potential throughput improvement. These tradeoffs indicate high potential for sparse gating.

**Figure 2:**
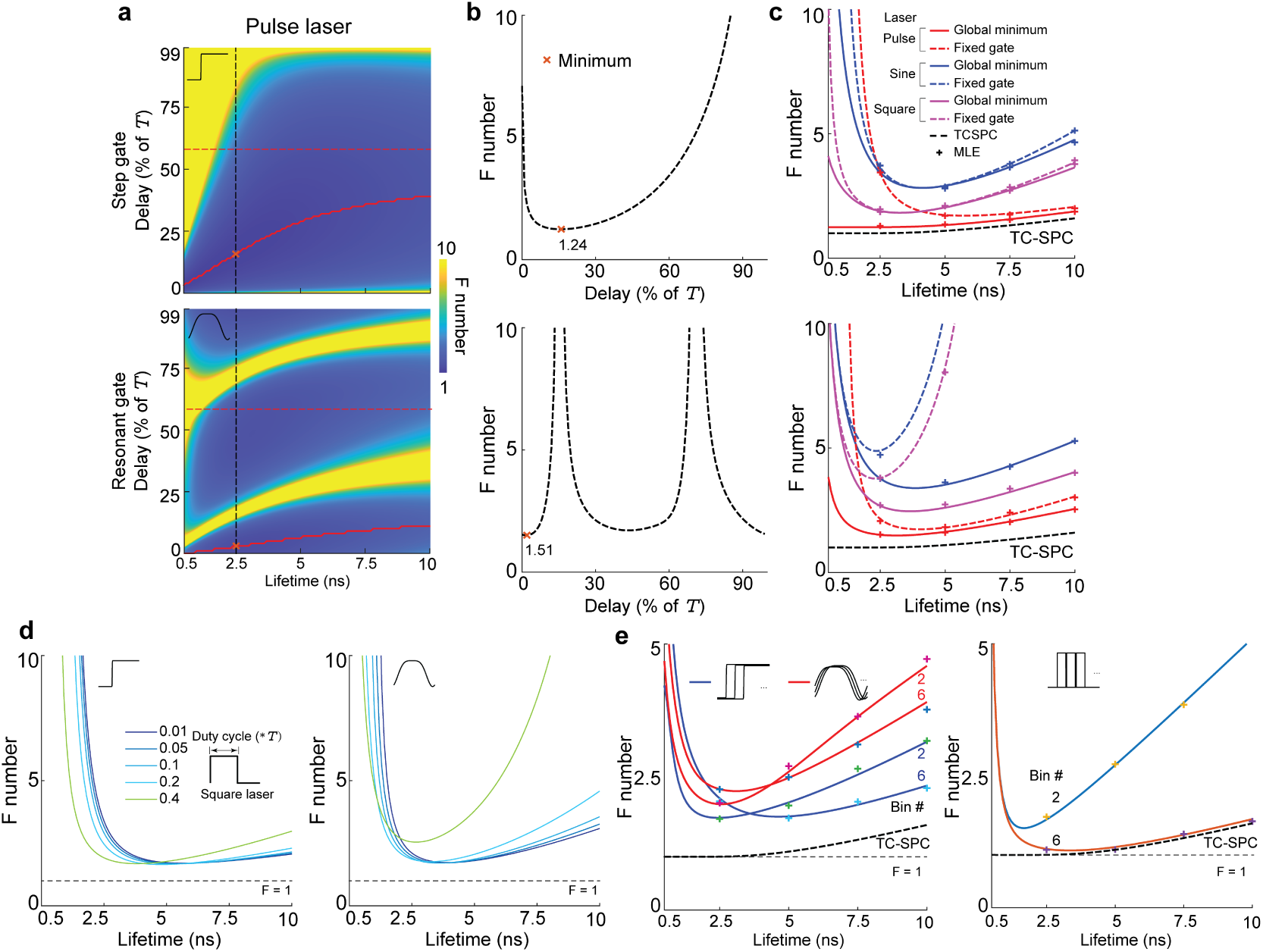
Single exponential estimation. **a)** Heatmaps (upper: step gate; lower: resonant gate) show F numbers as a function of lifetime *τ* and gate delay, calculated from CRLBs. The red dashed lines correspond to a gate delay of 0.6\**T* (laser repetition period *T* = 25 ns = 1 40 MHz). Red solid lines trace the minimum achievable F across all delays. The red dashed and solid lines are also plotted in panel (c). Black dashed lines show the F number with *τ* = 2.5 ns across delays, which are plotted in panel (b). The red cross indicates minimum achievable F for *τ* = 2.5 ns. **b)** Line cuts of F vs. gate delay (upper: step; lower: resonant) with *τ* = 2.5 ns corresponding to vertical lines in (a). Minimum F is indicated. **c)** Line cuts of F vs. *τ* for pulsed (red), sinusoidal (blue), or square (magenta) laser excitation waveforms. Dashed lines show the F number with a fixed gate delay (0.6\**T*, red dashed lines in (a)). Solid lines show the minimum achievable *F* across gate delays (solid red lines in (a)). The marked points show corresponding F number verified from Monte Carlo simulation and MLE. For comparison the black dashed lines shows the F number from TC-SPC (256 bins across the laser period). **d)** Illumination with square wave lasers of varying duty cycles can provide a cost-effective alternative for EO-FLIM excitation. F number with step gates (left) and resonant gates (right) are compared across duty cycles. **e)** Effect of increasing parallel (step / resonant) and series (square) gate number on single exponential estimation for a fixed 4 nanosecond spacing between gates and with photons equally distributed to each gate path. Data points show MLE verification for each line. Black dashed lines shows the F numbers from TC-SPC.

Unlike traditional TC-SPC approaches requiring a short pulsed laser, EO-FLIM can be applicable to different laser sources and temporally distributed excitation waveforms. This is a promising way to reduce system cost and achieve higher excitation powers, for example with modulated lasers. To simulate the impact of laser waveform on estimation, we next compared F numbers with different laser waveforms (Fig. 2c-d and Fig. S1). The penalty in F is approximately 1.7× (step) or 1.7× (resonant) going from pulsed to square wave illumination (50% duty cycle) and 3.1× (step) or 2.4× (resonant) to sine wave illumination (Fig. 2c and Fig. S1). For square wave illumination, the duty cycle can be tuned. We compared the F numbers across lifetimes with a fixed gate delay and with varying duty cycle. Increasing the duty cycle from 0.01 to 0.2 introduces little change of the F number across the lifetime range. Further increase of the duty cycle (up to 0.4) tends to decrease F numbers for short lifetimes and increase F numbers for long lifetimes (Fig. 2d). These results suggest that low cost lasers may find application for EO-FLIM and that optimal duty cycles can be chosen for a given experiment. In reality, perfectly sharp square waves are not achievable so we also tested the effect of rise time of square waveform illumination. F numbers increased with rise time. A 1-nanosecond 10–90% rise/fall time slightly increased the F numbers, introducing change factors of 1.3 for step or 1.2 for resonant gates compared to sharp edges (Fig. S2a-b). These indicate that we can effectively apply square excitation waveforms in EO-FLIM. Adding more gates in parallel requires dividing photons between gates, so it is not clear what the potential impact is on photon budget or design of EO-FLIM gating. To explore this, we analyzed F numbers using an increasing number of parallel EO-FLIM gates, with each gate having a fixed time step. We added gates with a 4 nanosecond time step with pulsed excitation of 25 ns period. Increasing the number of step or resonant gates can shift the lifetime range with minimal F number but has little impact on the F numbers themselves (Fig. 2e). This result is due to the trade-off between increased measurements and splitting of the photons as the number of gates increases. We also tested the performance of series square gates, approximating TC-SPC measurements with a limited number of wide bins. Increasing the number of square gates improved the estimation performance and the improvements plateaued at approximately 6 square gates with 4 ns bin width, spanning the whole 25 ns laser period. A 256-channel TC-SPC measurement only marginally improves F numbers, improving most significantly at short lifetimes where a 4 ns gate becomes too long to resolve the decay. (Fig. 2e).

In summary, we mapped the experimental space for EO-FLIM measurements of a single exponential. A single gate can approach shot noise limits of TC-SPC within a factor of 1.2 or 1.6. Going from pulsed to square or sine wave illumination comes with a penalty of 1.5 and 2.2 fold for step gates and 1.6 and 2.1 fold for resonant gates. Lastly, we found that increasing the number of step or resonant gates does not yield significant improvements for single exponential estimation. A set of 6 distributed gates with evenly divided photons can also perform well compared to ideal TC-SPC limits (1.05× for square, 1.53× for step, and 2.35× for resonant gates).

### 0.6 Double exponential decays with few gates

Double exponential decays such as those from FRET-based biosensors rely on deriving the lifetimes and fractions of the two lifetime components (e.g. substrate bound and unbound) to inform biology (25, 49). To evaluate optical gating for double exponential decays, we performed Fisher information analysis on EO-FLIM measurements starting with combinations of two and three gating waveforms.

There are four parameters (*τ*_1_, *N*_1_, *τ*_2_, *N*_2_) in a double exponential decay. Estimating all four unknown parameters is the most challenging case, requiring more than 2 gates and at least 4 independent measurements. However, in many cases with biosensors, we can gain prior knowledge of the decay by other means. For example we can measure the lifetime of the free donor fluorophore of a FRET pair, reducing the number of estimated parameters to 3. It will then require 2 gates with 4 measurements to estimate the double exponential decay. To explore this, we analyzed the CRLBs and the corresponding F numbers with one known lifetime component within the 2-gate combination delay space. The results showed that with prior known *τ*_2_, two gates are sufficient to estimate the rest of the parameters (*τ*_1_, *N*_1_, *N*_2_) with both step and resonant gates (Fig. 3a-h). Step gates perform better than resonant gates (by a factor of 1.8). From sweeping the 2-gate delay space, we can find the optimal 2-gate combination that enables the best estimation performance (Fig 3a-b and 3e-f). Using this fixed combination of two gates, we can further explore the parameter space by varying *τ*_2_ *τ*_1_ and *N*_2_ *N*_1_ ratios (Fig. 3c-d and 3g-h). As expected, when varying *τ*_2_ *τ*_1_ ratios, the F numbers increase drastically as *τ*_1_ and *τ*_2_ become closer to each other both in EO-FLIM and TC-SPC. The performance of two gate EO-FLIM with one known lifetime component approaches TC-SPC performance (Fig. 3d and 3h) by around 2.9-fold for resonant gates and 1.6-fold for step gates.

**Figure 3:**
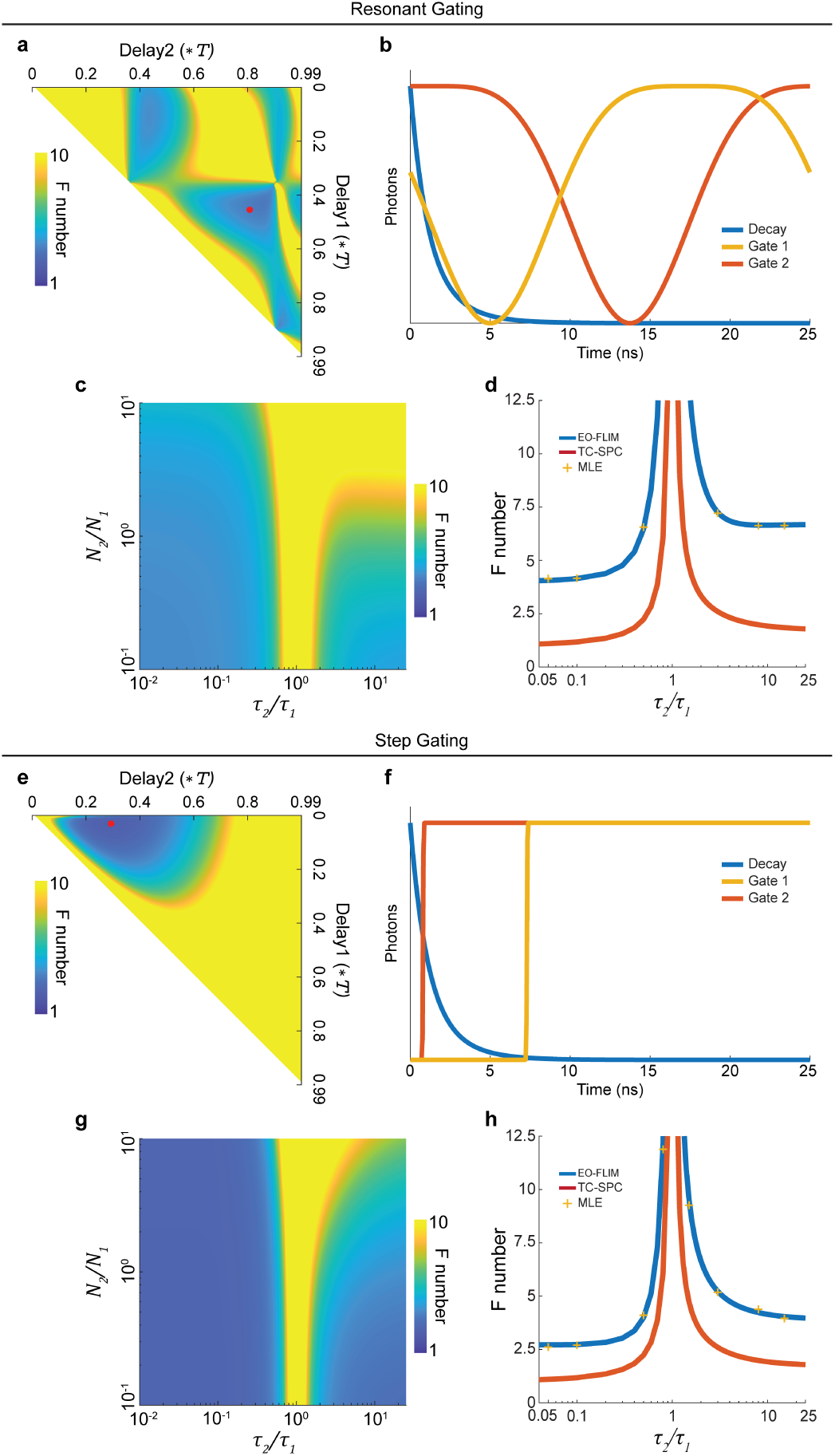
Estimating double exponential decays with one known lifetime using two complementary gates. **a)** F number of *τ*_1_ estimation as a function of two resonant gate delays on a double exponential decay with known *τ*_2_ (*τ*_1_ = 2 ns, *τ*_2_ = 1 ns, *N*_1_ = 10^7^, *N*_2_ = 10^7^). The two gates are in parallel meaning each gate samples half of the photons available to the measurement. The red dot indicates one of the delay combinations that give low F number, which is plotted in (b). **b)** Two optimal resonant gate functions. **c)** Using this fixed delay setting, we can then map F number for *τ*_1_ estimation with different combinations of *τ*_1_, *τ*_2_, *N*_1_, and *N*_2_ (*τ*_1_ = 2 ns, *N*_1_ = 10^7^). **d)** Comparison of F numbers for *τ*_1_ estimation with resonant gates to TC-SPC. The EO-FLIM line is generated using the gates from (b) at every *τ*_2_/*τ*_1_ ratio with *N*_1_ = *N*_2_. Data points represent MLE verification on Monte Carlo simulated data. **e)** F number of *τ*_1_ estimation as a function of two step gate delays, on the same double exponential decay as (a). The red dot indicates the optimal combination plotted in (f). **f)** Two optimal step gate functions. **g)** F number for *τ*_1_ with different combinations of *τ*_1_, *τ*_2_, *N*_1_, and *N*_2_, at the fixed step gate delays in (f). **h)** Comparison of the F numbers with step gates from (f) for EO-FLIM and TC-SPC with data points from MLE verification (*N*_1_ = *N*_2_).

We also explored the feasibility of deploying multiple gates to measure the full parameters of bi-exponential decays. Without prior knowledge of the decay, all four parameters (*N*_1_, *τ*_1_, *N*_2_, *τ*_2_) must be estimated, which requires a Fisher information matrix of rank four and four linearly independent measurements. In EO-FLIM, however, each complementary pair satisfies *λ_G_ λ_U_* = *N*_1_ *N*_2_, so k parallel gates provide 2k measurements but only k + 1 independent ones: the shared total intensity plus one gated fraction per gate. Therefore in the case of parallel gates one gate is sufficient for single exponential estimation, two gates are needed for a bi-exponential decay with one known lifetime, and at least three gates (six measurements) are required for a fully unknown bi-exponential decay. To explore the performance for double exponential estimation, we analyzed the 3-gate combination delay parameter space (Fig. S3). The results showed that 3 gates are mathematically sufficient to estimate the 4 parameters of the double exponential decays (*τ*_1_ = 2 ns, *τ*_2_ = 1 ns, *N*_1_ = 10^7^, *N*_2_ = 10^7^) with both step and resonant gates. The performance of step gates are much better than resonant gates (Fig. S3a-f). From this analysis, we can also find the optimal 3-gate combinations that enable the best performance. The best gate combinations for *τ*_1_ or *τ*_2_ estimation are not identical but share similar gate delays (Fig. S3a and S3d). The resulting best gating schemes for estimating both *τ*_1_ and *τ*_2_ are (0.41, 0.13, 0.01)\**T* for step gates and (0.93, 0.90, 0.41)\**T* for resonant gates. These gates capture the maximum information by optimally interacting with the decay (Fig. S3b and S3e). We can also find the best gating schemes across different *τ*_1_ and *τ*_2_ combinations (Fig. S3c and S3f). As with all double exponential cases, F number diverges as the lifetime components *τ*_1_ and *τ*_2_ become closely spaced. This case is the most challenging, and gating approaches here have a significant penalty relative to TC-SPC, especially with resonant gates (2.5 with three step and 6.7 with three resonant gates for *τ*_1_ estimation in the case analyzed). However the throughput advantage of optical gating could still compensate for the information loss in applications with large photon numbers, and adding more gates can improve performance (following section).

### 0.7 Double exponential decays with an increasing number of gates

While three gates are theoretically sufficient for estimating double exponential decays, sparse gated measurements give up significant information relative to TC-SPC. We next analyzed whether increasing the number of parallel gates while splitting photons evenly between each measurement could improve performance. We analyzed increasing numbers of parallel step and resonant as well as corresponding numbers of series square gates. We used both even gate spacing in time within the laser repetition period or fixed 1 ns intervals between gates on decays with (Fig. 4) or without (Fig. S4) prior knowledge of one lifetime parameter.

**Figure 4:**
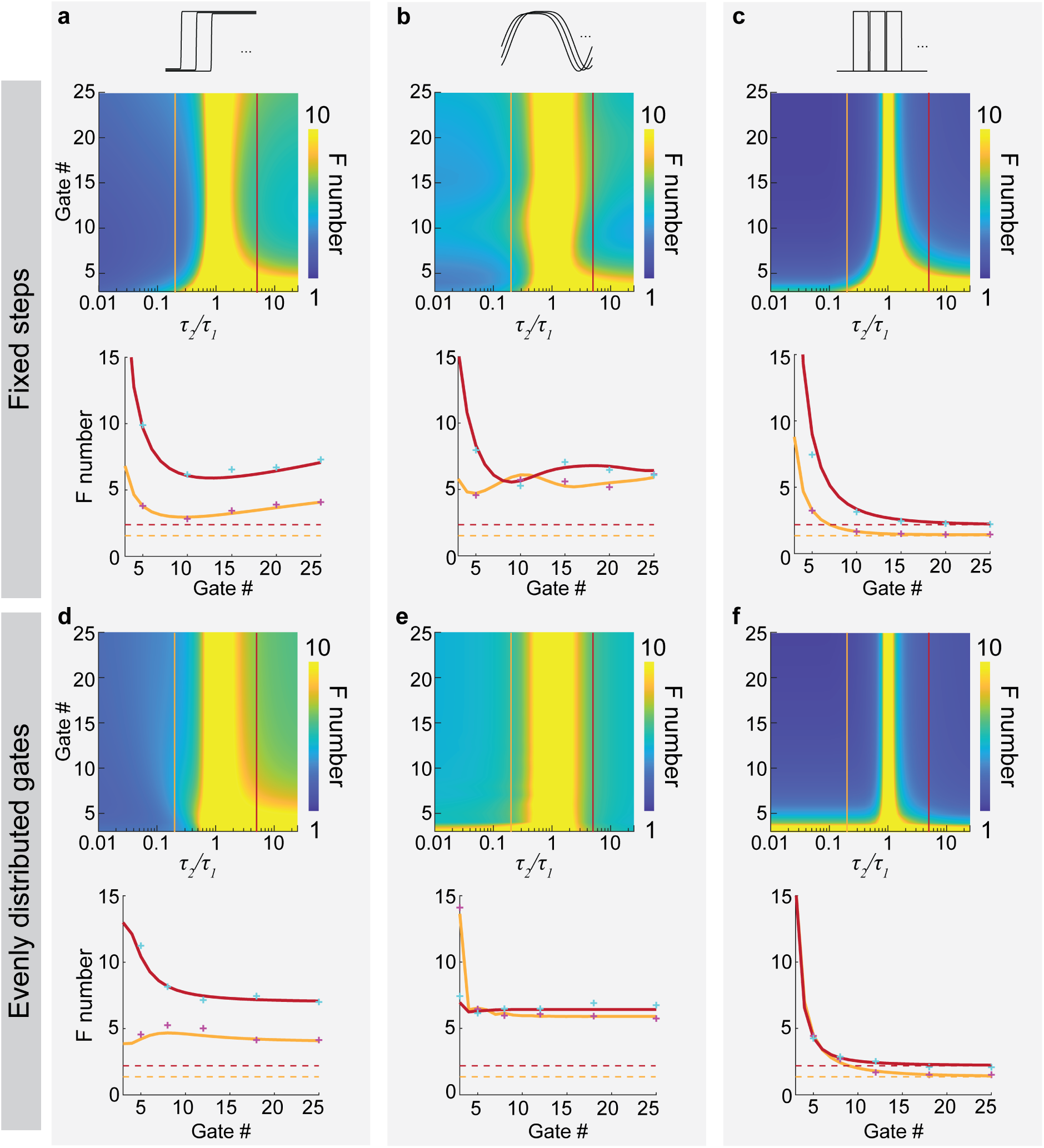
Estimating double exponential decays with one known lifetime using many parallel or series gates. **a-f)** Heatmaps (upper panels) and line plots (lower panels) showing F numbers for *τ*_1_ estimation, with increasing number of parallel step (a and d), resonant (b and e), and series square (c and f) gate functions. The gates are arranged within the excitation period (25 ns) at fixed 1 ns intervals (a-c) or evenly distributed (d-f). Line plots show F numbers as gate number is increased at *τ*_2_/*τ*_1_ = 0.2 or 5, corresponding to the lines in the heatmaps (*τ*_1_ = 2 ns, *N*_1_ = *N*_2_ = 107). Dashed lines are the TC-SPC F numbers for *τ*_1_ with corresponding parameter conditions. Indicated data points provide MLE verification on Monte Carlo simulated data. It is noteworthy that across all panels performance converges within 5-10 gates and that going slightly beyond the minimal case of 3 gates (Fig. S3) recovers significant information.

With *τ*_2_ known, estimation precision for both fixed and evenly distributed gates asymptotically converged for *τ*_1_ estimation (Fig. 4). For step gates, increasing the number of gates reached asymptotic performance after about 10 gates at both low and high *τ*_2_ *τ*_1_ ratios (Fig. 4a and d). Resonant gates have more oscillatory behavior due to how the gate edges fall relative to the decay (Fig. 4b). For resonant gates, the performance notably converged with fewer number of gates compared to step gates (Fig. 4b and e). For example, with evenly spaced gates, the F number asymptotes rapidly after 4 gates at low *τ*_2_ *τ*_1_ ratios and stayed constant across gate numbers whereas at high *τ*_2_ *τ*_1_ ratio, only 3 gates reaches the asymptote (Fig. 4e). For square gates, emulating TC-SPC, we approach the infinite bin limit after about 10 gates at both low and high *τ*_2_ *τ*_1_ ratios with both evenly distributed and fixed spaced gates (Fig. 4c and f).

Without prior knowledge of the decay, the F numbers for *τ*_1_ estimation also showed similar convergence and asymptotic behaviors as gate number is increased (Fig. S4). As expected, performance is worse than the case with *τ*_2_ known, especially for fixed interval gating. Step gates with fixed intervals only reached the asymptotes after about 20 gates at high *τ*_2_/*τ*_1_ ratio, while around 10-12 gates are still required at low *τ*_2_/*τ*_1_ ratio or with evenly distributed gates at both low and high *τ*_2_/*τ*_1_ ratio (Fig. S4a and S4d). Resonant gates required 15 or 4 gates to reach the asymptotes in the fixed or evenly distributed gate cases at low *τ*_2_/*τ*_1_ ratio (Fig. S4b and S4e). For square gates, gates with fixed steps required more than 15 gates to reach the asymptotes, compared to 12 gates for evenly distributed ones (Fig. S4c and S4f).

In summary, we see convergence across gating waveforms and delay spacings with increasing gate number. For one known lifetime, convergence is faster (5-10 gates) than the case with no prior information (10-15 gates). It is also notable that resonant gates converge especially fast and give stable estimation across gate numbers, although they asymptote to a higher F number overall. These results indicate potential designs for high-throughput multiplexing leveraging optical gates.

## DISCUSSION

FLIM imaging is emerging as a powerful tool across neuroscience and biological measurement. Developing fluorescence lifetime biosensors enable readout of signals in live tissue and behaving organisms, ranging from voltage (2, 50–54) and calcium indicators (55–57) for fast neural activity to sensors reporting slower dynamics such as neuromodulators (3, 4, 58, 59), intracellular signaling (7–9, 49, 60–62), and metabolites (5, 6). Fluorescence lifetime imaging also provides an orthogonal dimension for probe multiplexing in addition to wavelength (29–31). Applications of FLIM, however, have been limited by low photon throughput sensors and point-scanning measurements. In fact, for many applications imaging biosensors existing FLIM techniques do not take advantage of the full photon flux available to intensity measurements (2, 22). Optical gating provides an approach to perform FLIM measurements on low-noise intensity camera sensors in wide-field, improving throughput and dynamic range.

In this study, we provide a comprehensive analysis of gating-based FLIM measurements using Fisher information theory. We focus on new classes of lossless gate waveforms that may be realized all-optically in EO-FLIM. We explored lifetime estimation performance for both single and dual exponential decays and analyzed step, square, and resonant gate waveforms along with pulsed, square, and sinusoidal laser illumination. To independently verify our results we used Monte Carlo simulation and maximum likelihood estimation. We found four summary conclusions:

1. A single gate is sufficient for single exponential estimation within 1.2-1.6 TC-SPC limits. There is a large degree of flexibility in the usable parameters for excitation and gating waveforms, allowing diverse and potentially low-cost FLIM implementations to be designed.
2. Two or three gates can be optimally chosen to estimate double exponential decays with or without prior information of one lifetime component. Fisher information informs the choice of optimal gate delays and allows estimation 1.6 / 2.9 fold of the TC-SPC limits for one known (two step / resonant gates) and 2.5 / 6.7 fold with no prior information (three step / resonant gates).
3. Large numbers of gates are not necessarily needed for efficient bi-exponential lifetime estimation. As more than three gates are added, performance becomes more robust when estimating double exponential decay components over larger parameter spaces. With prior knowledge of one lifetime component, only 5-10 distributed step or square gates are required. Resonant gate waveforms can also perform robustly across parameters and only require 3-4 gates to converge to asymptotic limits.
4. In the case of no prior information (i.e. four unknowns) it is challenging to estimate lifetimes using resonant gates, incurring an F ratio of > 8.8 compared to TC-SPC. Using step function gating, however, recovers performance to 3.5 fold of TC-SPC limits, while square gating converges to the TC-SPC limit within 10-15 bins.

We note that in many experimental contexts with biosensors, prior information about a probe or biosensor may be inferred from spatial and/or temporal context in the dataset, or from prior probe calibration experiments. All analysis here ignores spatial and temporal context and focuses on blind unmixing of decay dynamics from a single image pixel, aggregate region of interest, or fiber photometry output. We also note that complete parameter estimation is a distinct problem from achieving probe separability, so in applications that only require multiplexing a known basis of fluorescent probe signatures, the requirements on photon number may be dramatically reduced (38).

A key distinction of the EO-FLIM approaches analyzed here is that they capture time information by gating before the photon detection event, separating the FLIM measurement architecture from electronic detector noise. Many existing approaches introduce excess experimental noise. For example, direct digitization of a lifetime decay can speed up point-scanning measurements by an order of magnitude but can also introduce additional noise from the photodetector, analog-to-digital conversion, and amplification (63, 64). We only consider shot noise because optical gating can use low noise intensity detectors. This is justified because scientific camera sensors routinely achieve pixel read noise below 1 electron per pixel. We also restrict ourselves to ideal gating waveforms and note that in practice there may be incomplete modulation depth and wavelength-dependent gating. Our FI simulation framework can be adapted to these cases. We note that our MLE results are in the limit of large N where we expect estimation to be asymptotically unbiased. This limit is justified for applications imaging bright fluorescent biosensors or densely-expressed probes from populations of cells in the brain, for example in fiber photometry. Our results generalize to any gating method that achieves lossless gating, making them relevant not only for EO-FLIM but also for next generation in-camera gating architectures based on SPAD arrays or multi-tap pixels that are becoming possible with improved semiconductor process (65–67).

## CONCLUSION

Our information theory framework informs the optimal design of next-generation gated FLIM measurements to maximize per-photon information. Conventional FLIM techniques have taken the approach that more time bins is better. We argue that a few optimal bins can be equally, if not more informative after taking into account potential gains in photon throughput and decreased measurement noise afforded by optical gating. In many scenarios simple experiments can be designed using a few optical gates to achieve lifetime estimation performance close to fundamental limits.

## DATA AND CODE AVAILABILITY

Example code will be made available. Simulated data and analysis codes are available from the authors upon request.

## AUTHOR CONTRIBUTIONS

P.M. and A.B. conceived the study. All authors contributed to simulation development and manuscript revision.

## ACKNOWLEDGMENTS

This work was supported by funding from NIH R35 GM166021, the Biohub Imaging Residency Program, the Dan and Martina Lewis Biophotonics Fellowship (L.H.), the Pioneer Fund Postdoctoral Scholar Award (P.M.), and the Kavli Institute for Brain and Mind Postdoctoral Scholars Program (P.M.).

## DECLARATION OF INTERESTS

A.J.B. is an inventor on patents PCT/US2019/062640; US 11,965,780; and US 11,828,851 describing wide-field, scanning, and resonant EO-FLIM.

## DECLARATION OF GENERATIVE AI

During the preparation of this work the authors used Claude and Claude Code by Anthropic in order to develop and check simulation codes and derivations. The authors reviewed and edited all outputs and take full responsibility for the content of the publication.

## SUPPLEMENTARY FIGURES

**Figure S1:**
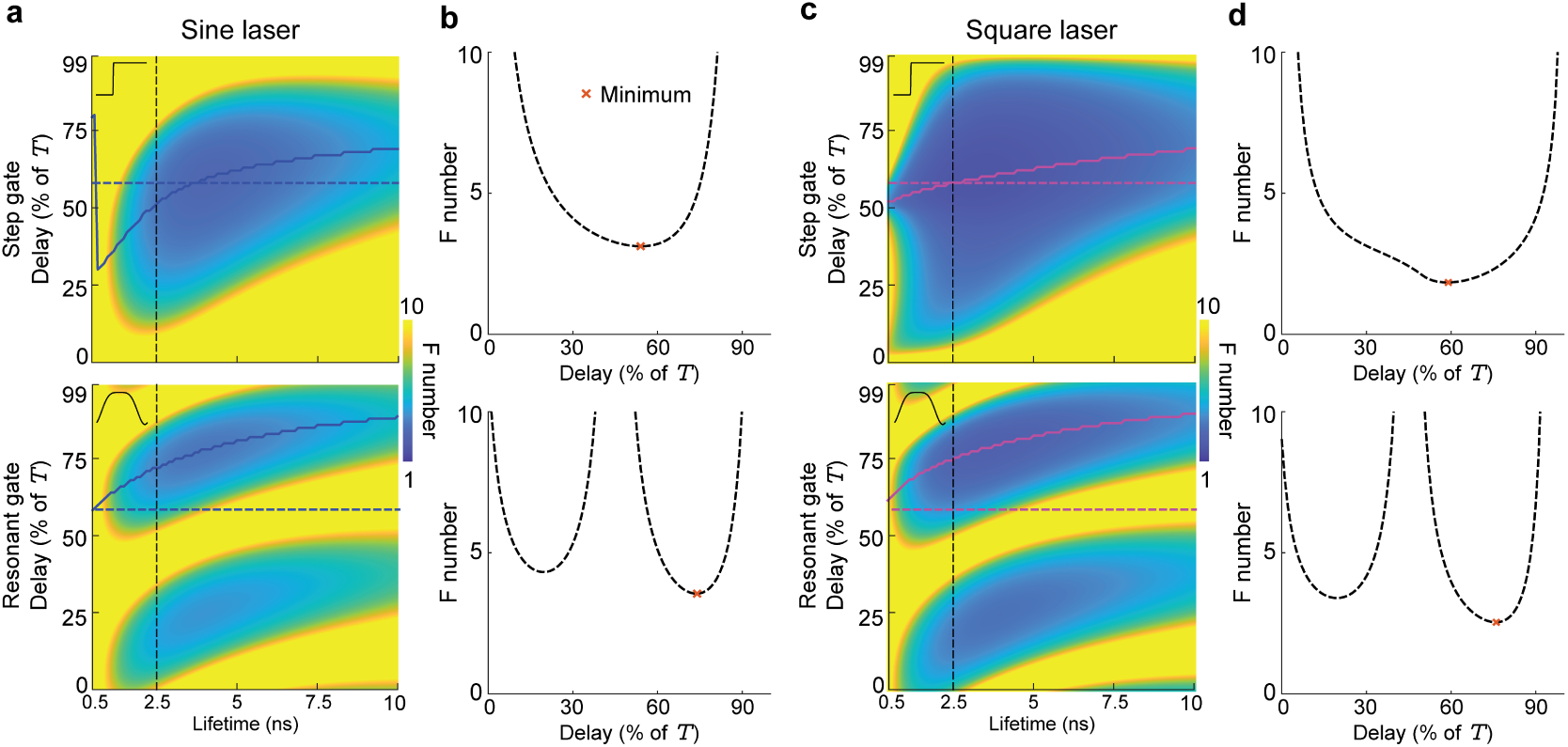
Single exponential estimation with sinusoidal and square laser waveforms. **a)** Heatmaps (upper: step gate; lower: resonant gate) show F numbers as a function of lifetime *τ* and gate delay, calculated from CRLBs, with sinusoidal laser excitation. The blue dashed lines correspond to a gate delay of 0.6\**T*. Blue solid lines trace the minimum achievable F across all delays. The blue dashed and solid lines are also plotted in Fig. 2c. Black dashed lines show the F number with *τ* = 2.5 ns across delays, which are plotted in panel (b). **b)** Line cuts of F vs. gate delay (upper: step; lower: resonant) with *τ* = 2.5 ns corresponding to vertical lines in (a). Minimum F is indicated. **c)** Heatmaps (upper: step gate; lower: resonant gate) show F numbers as a function of lifetime *τ* and gate delay, calculated from CRLBs, with square laser waveform. The magenta dashed lines correspond to a gate delay of 0.6\**T*. Magenta solid lines trace the minimum achievable F across all delays. The magenta dashed and solid lines are also plotted in Fig. 2c. Black dashed lines show the F number with *τ* = 2.5 ns across delays, which are plotted in panel (d). **d)** Line cuts of F vs. gate delay (upper: step; lower: resonant) with *τ* = 2.5 ns corresponding to vertical lines in (c). Minimum F is indicated.

**Figure S2:**
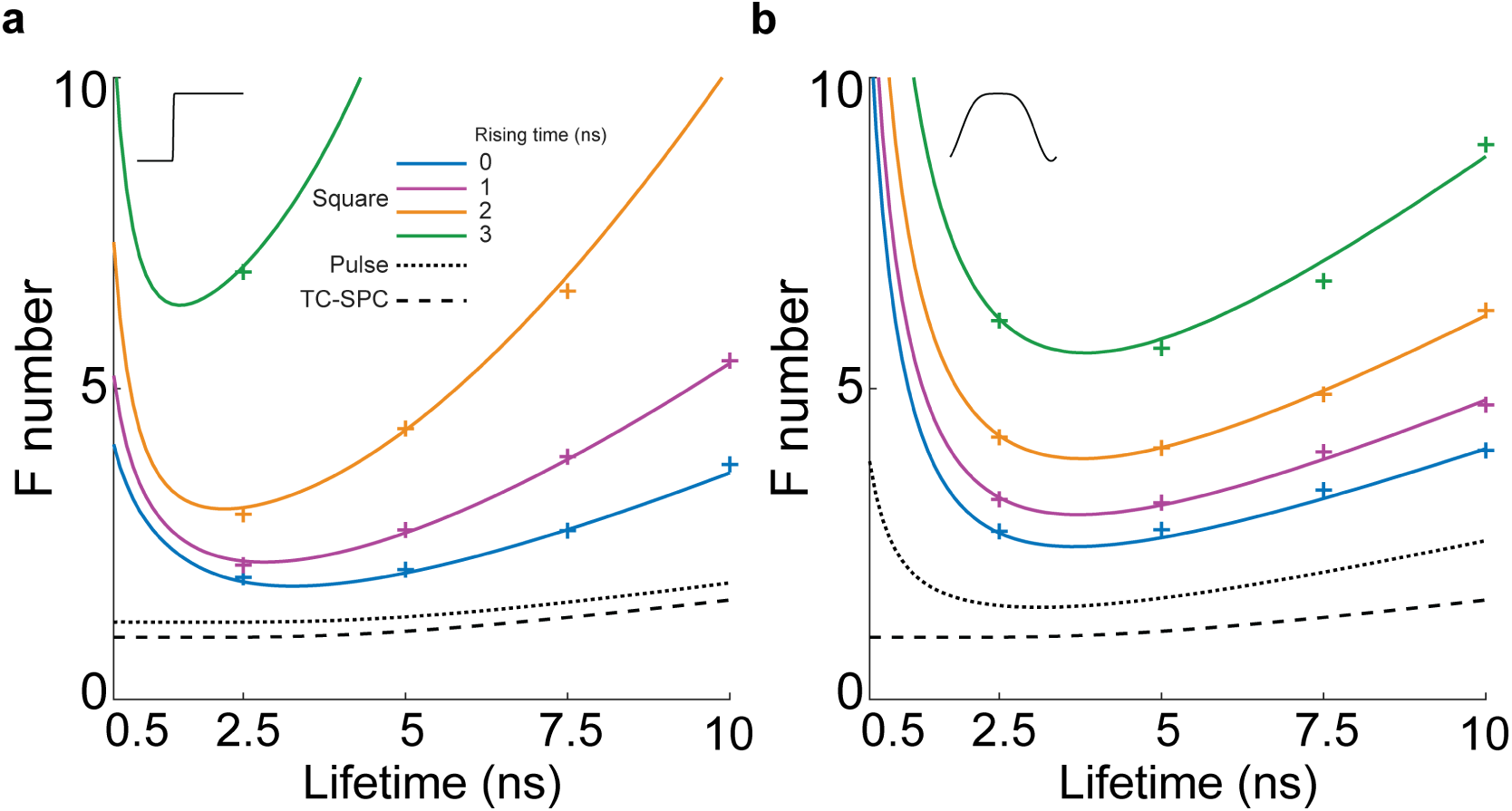
Effects of square wave excitation rise / fall time. **a-b)** Minimum F numbers (at optimal gate delays) for a single exponential using square laser waveforms (duty cycle = 0.5) with different 10-90*%* rise / fall times, by step (a) or resonant (b) gates in EO-FLIM. Dashed lines show the minimum F numbers achieved with a pulsed laser in EO-FLIM and TC-SPC.

**Figure S3:**
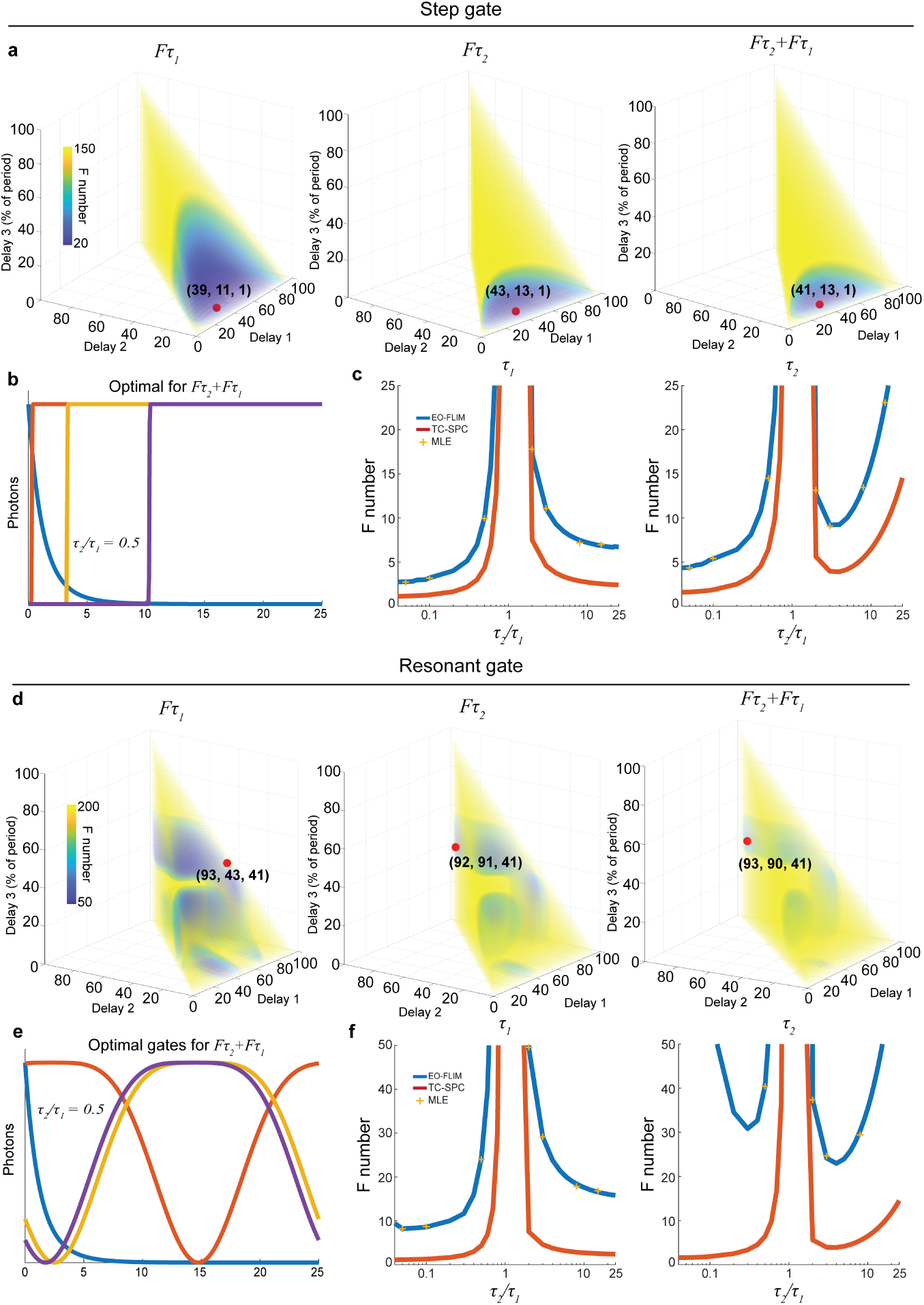
Estimating double exponential decays using three gates and no prior information. **a)** Heatmaps showing the F numbers of *τ*_1_, *τ*_2_, and summed *τ*_1_ and *τ*_2_ estimation as a function of three step gate delays on a double exponential decay with no prior information (*τ*_1_ = 2 ns, *τ*_2_ = 1 ns, *N*_1_ = 10^7^, *N*_2_ = 10^7^). The three gates are in parallel meaning each gate samples one third of the photons available to the measurement. The red dots indicate one of the optimal delay combinations. **b)** Plots of double exponential decay and the corresponding three-gate combination that gives the minimum sum of the F numbers for both *τ*_1_ and *τ*_2_ estimation, indicated as the red dot in (a, right). **c)** F number for *τ*_1_ or *τ*_2_ with the optimal combination of the three step gates from (b) on decays with different *τ*_1_ and *τ*_2_ combinations (*τ*_1_ = 2 ns, *N*_1_ = *N*_2_ = 10^7^). **d)** Heatmaps showing the F numbers of *τ*_1_, *τ*_2_, and summed *τ*_1_ and *τ*_2_ estimation as a function of three resonant gate delays on a double exponential decay with no prior information of the decay parameters (*τ*_1_ = 2 ns, *τ*_2_ = 1 ns, *N*_1_ = 10^7^, *N*_2_ = 10^7^). The red dots indicate optimal delay combinations. **e)** Plots of double exponential decay and the corresponding three-gate combination that gives the minimum sum of the F numbers for both *τ*_1_ and *τ*_2_ estimation, indicated as the red dot in (d, right). **f)** F number for *τ*_1_ or *τ*_2_ with the optimal combination of three resonant gates from (e) on decays with different *τ*_1_ and *τ*_2_ combinations (*τ*_1_ = 2 ns, *N*_1_ = *N*_2_ = 10^7^).

**Figure S4:**
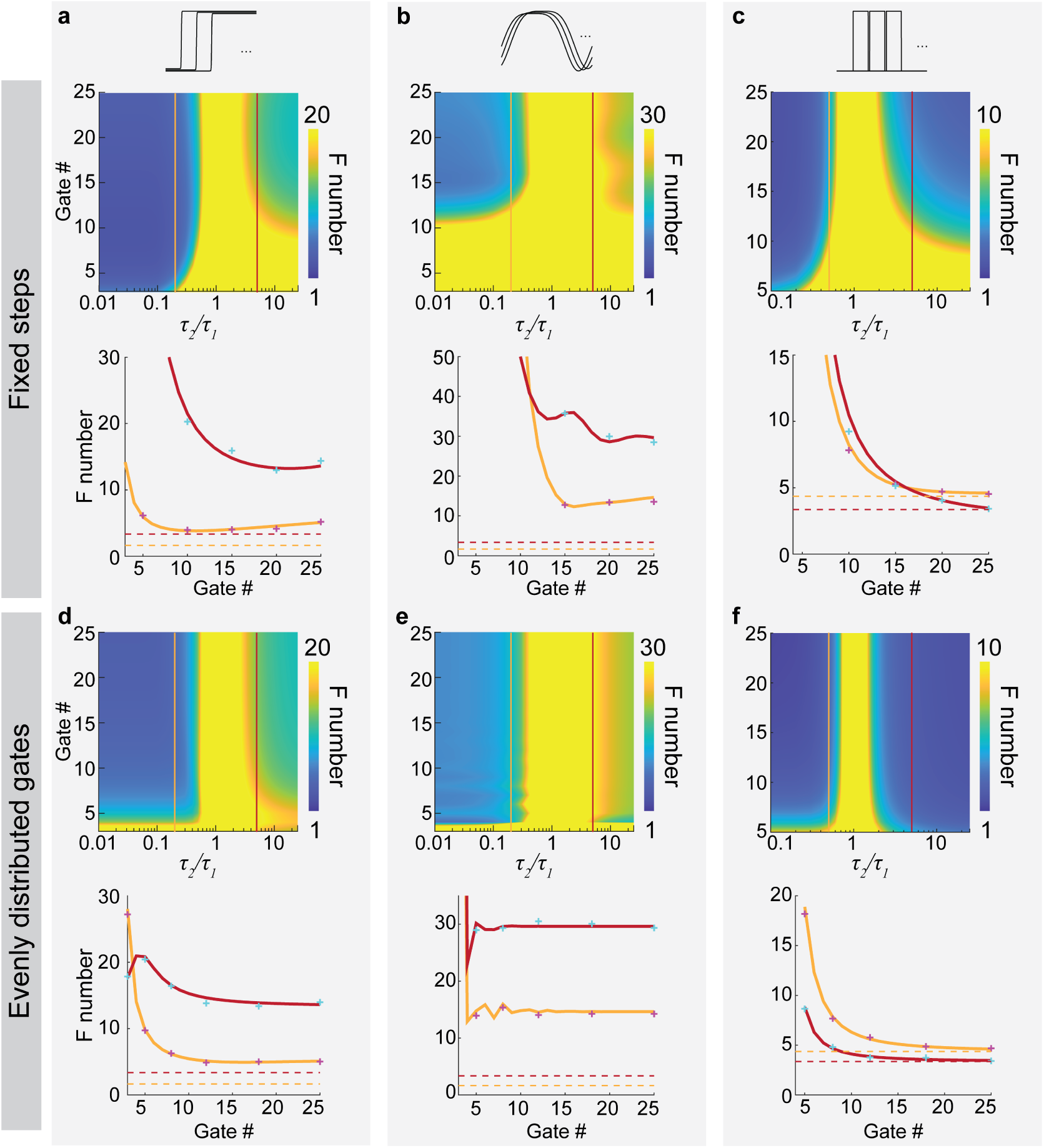
Estimating double exponential decays using many gates without prior information. **a-f)** Heatmaps (upper panels) and line plots (lower panels) showing F number for *τ*_1_ estimation, with increasing number of parallel step (a and d), resonant (b and e), and series square (c and f) gate functions. The gates are arranged within the excitation period (25 ns) at fixed 1 ns intervals (a-c) or evenly distributed (d-f). Line plots show F number as number of gates is increased at *τ*_2_/*τ*_1_ = 0.2 or 5 (a-b and d-e) or 0.5 and 5 (c and f), corresponding to the lines in the heatmaps (*τ*_1_ = 2 ns, *N*_1_ = *N*_2_ = 10^7^). Dashed lines are the TC-SPC F numbers for *τ*_1_ with the same estimation conditions. Indicated data points provide MLE verification from Monte Carlo simulated data. For square gates here we swept parameters using a larger gate number and *τ*_2_*/τ*_1_ ratio (5 gates and ratio = 0.1 instead of 3 gates and ratio = 0.01), because smaller numbers of square gated measurements were not able to resolve small *τ*. It is noteworthy that across all panels performance converges within 10-15 gates, except in the top right panel where fixed spacing requires 15-20 gates.

### Section S1 FISHER INFORMATION ANALYSIS FOR EO-FLIM

#### Section S1.1 Fluorescence decay under different excitation waveforms

For every excitation waveform, the photon probability in bin *t_k_* is obtained by normalizing the steady-state fluorescence *F_i_*(*t*) over one period, and its lifetime derivative follows from that of *F_i_*:

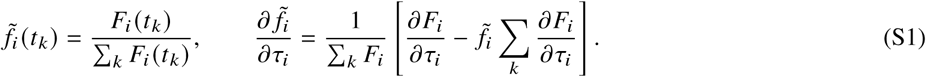

For pulsed excitation this is identical to the bin integral in the Methods. Below we give *F_i_*(*t*) and *∂F_i_*/*∂τ_i_* for each waveform, with 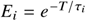.

##### Pulsed excitation

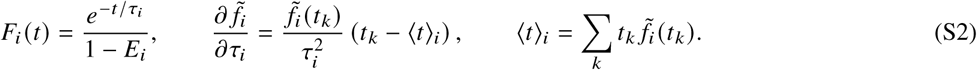

##### Sinusoidal excitation

For 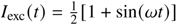 with *ω* = 2*π*/*T*:

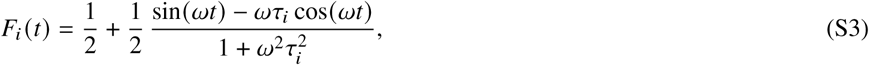

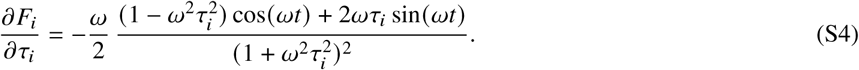

##### Square-wave excitation

The laser is on for 0 ≤ *t* < *t*_on_ = *dT* (duty cycle *d*) and off for *t*_off_ = *T* − *t*_on_. With 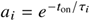 and 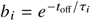:

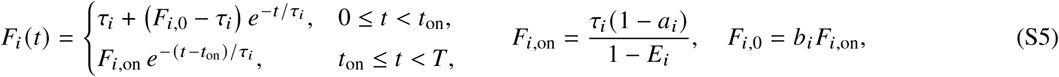

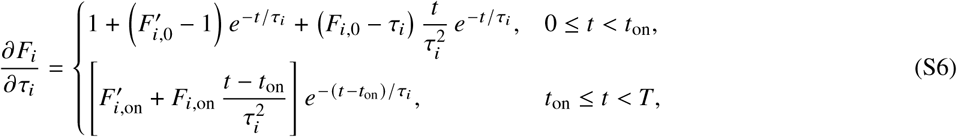

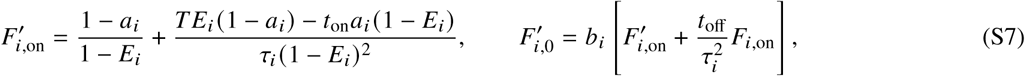

where primes denote *∂*/∂*τ_i_*.

##### Square wave with finite edges

Each laser edge is a logistic function with 10–90% transition time Δ*t_L_*:

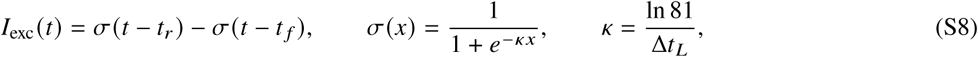

with *t_r_* = *w*/2 and *t_f_* = *t*_on_ + 3*w*/2, where *w* = Δ*t_L_* ln 99/ln 9 ≈ 2.09 Δ*t_L_* is the 1–99% width of each edge. This places *t* = 0 at the 1% point of the rising edge and keeps a flat top of duration *t*_on_. *I*_exc_(*t*) is wrapped periodically over *T*. *F_i_* is computed as the periodic solution of

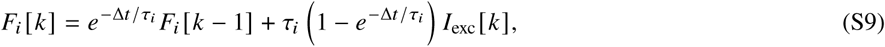

and ∂*F_i_*/∂*τ_i_* is computed numerically.

#### Section S1.2 Gate functions

Each gate *g_m_*(*t*) (0 ≤ *g_m_* ≤ 1) has a complementary ungated channel 1 − *g_m_*(*t*).

##### Step gates

A logistic step *s_m_*(*t*) with gate duty cycle *d_g_*, delay ϕ*_m_T* and 10–90% rise time Δ*t_r_* is converted to the optical transmission

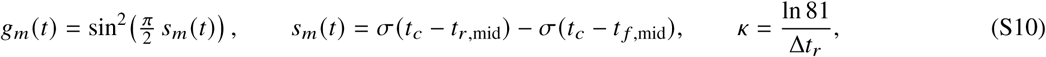

where *t_c_* = mod(*t* − ϕ*_m_T*, *T*) and *t_r_*_,mid_, *t_f_* _,mid_ are the edge midpoints. For all sharp step gates analyzed here, we apply a 100 picosecond 10–90% rise time and the gate extends until the end of the laser period.

##### Resonant gates

For a modulator driven at *f_g_* = *f*_rep_/2:

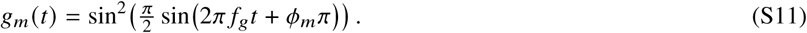

Note that this functional form results from the transmission of light through crossed polarizers after undergoing a birefringent phase shift in a resonantly driven Pockels cell or electro-optic modulator.

#### Section S1.3 Fisher information matrix

The elements of the symmetric matrix *I* are defined below, summing over all 2*M* measurements:

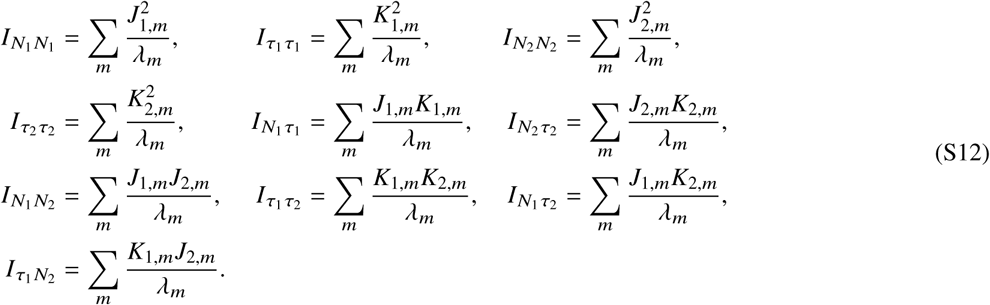

#### Section S1.4 Temporal Fisher information density

The information per unit time on parameter *θ* (Fig. 1d) is

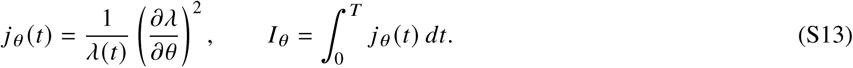

For a bi-exponential decay, *λ*(*t*) = Σ*_i_N_i_p_i_* (*t*) with 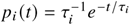, this gives

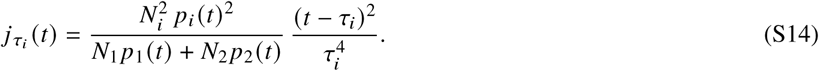

For a single component, *j_τ_* (*t*) = *N* (*t* − *τ*)^2^*e*^−*t*/*τ*^/*τ*^5^ and *I_τ_* = *N*/*τ*^2^, which corresponds to *F* = 1.

### Section S2 FISHER INFORMATION FOR TCSPC

Following Bouchet *et al.*[32], with *u_i_* = *i*Δ*t*/*τ*_1_, *r* = *T*/*τ*_1_ and *γ* = *τ*_1_/*τ*_2_, the bin probabilities and their logarithmic lifetime derivatives are

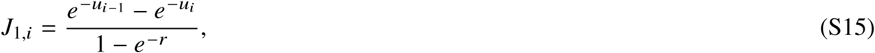

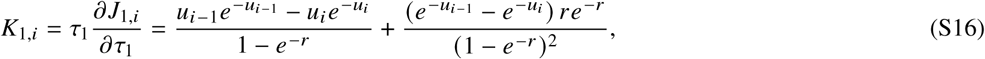

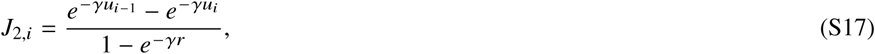

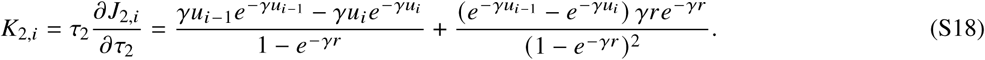

With *λ_i_* = *N*_1_*J*_1,_*_i_* + *N*_2_*J*_2,_*_i_*, the Fisher information matrix elements are

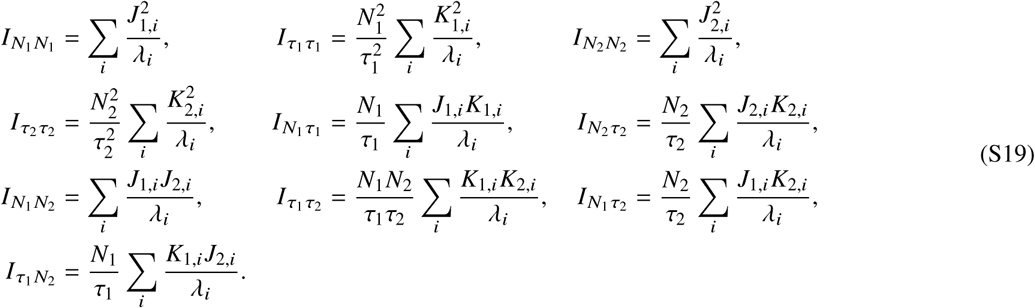

Note that our TC-SPC model does not consider finite excitation pulse width, timing jitter, or pile-up effects, which can all degrade estimation performance in real experiments[28].

#### Section S3 MAXIMUM LIKELIHOOD ESTIMATION AND MONTE CARLO SIMULATION

The log-likelihood in the Methods was evaluated with 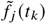 from the same excitation waveform used to simulate the data. The duty cycle and laser transition time were fixed at their known values. The parameters were estimated by minimizing −ℓ(***θ***) with MATLAB’s fmincon, with *τ_j_* and *N _j_* restricted to fixed ranges. For TC-SPC, the optimization was performed over ln *N _j_* to improve numerical conditioning.

For each parameter set, 1000 datasets were drawn from Poisson distributions with means *λ_G,m_* and *λ_U,m_* (or *λ_i_* for TC-SPC) and fitted by MLE. The empirical F number,

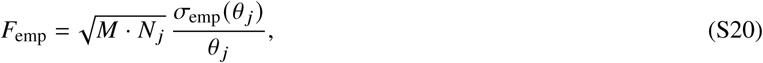

where *σ*_emp_ is the standard deviation of the estimates across trials, which was then compared with the value predicted from the CRLB. M accounts for equal photon division among parallel gates (step and resonant). For series square gating M = 1 here.

